# Semi-automated analysis of beading in degenerating axons

**DOI:** 10.1101/2025.02.05.636573

**Authors:** V C Pretheesh Kumar, Pramod Pullarkat

## Abstract

Axonal beading is a key morphological indicator of axonal degeneration, which plays a significant role in various neurodegenerative diseases and drug-induced neuropathies. Quantification of axonal susceptibility to beading using neuronal cell culture can be used as a facile assay to evaluate induced degenerative conditions, and thus aid in understanding mechanisms of beading and in drug development. Manual analysis of axonal beading for large datasets is labor-intensive and prone to subjectivity, limiting the reproducibility of results. To address these challenges, we developed a semi-automated Python-based tool to track axonal beading in time-lapse microscopy images. The software significantly reduces human effort by detecting the onset of axonal swelling. Our method is based on classical image processing techniques rather than an AI approach. This provides interpretable results while allowing the extraction of additional quantitative data, such as bead density, coarsening dynamics, and morphological changes over time. Comparison of results obtained through human analysis and the software shows strong agreement. The code can be easily extended to analyze diameter information of ridge-like structures in branched networks of rivers, road networks, blood vessels, etc.

## 1 Introduction

Axons are thin tubular extensions of neuronal cells which help in the conduction of electrical signals over long distances. When neurons undergo degeneration as a result of disease, injury or ageing, axons develop strong modulations of thickness and eventually lose their ability to function [1, 2, 3, 4, 5, 6]. Axonal beading, which is a morphological transformation of axons resulting in a modulation of axonal diameter, and axonal retraction, are commonly observed shape changes which are indicative of damage to axonal internal structure [7]. Interestingly, there have been very recent reports of normal axons exhibiting a beaded geometry in brain slices or in whole organisms [8, 9, 10].

Morphological transformations of axons can be studied in the laboratory using neuronal cell culture and the resulting shape evolution can be quantified [7]. Such studies are important in understanding normal axonal function [11, 12], as well as the diseased conditions mentioned above. Investigations on morphological changes are also important in investigating the mechanisms of axonal beading caused by cytoskeletal damage. Also, quantification of axonal beading can be used as a facile assay to screen neuropathy causing anti-cancer drugs for their potential to cause axonal damage, apart from improving our understanding of axonal degeneration process.

Our laboratory has extensively studied the impacts of anti-cancer drugs on chick dorsal root ganglia neurons. For this we quantify the evolution of axonal beading by measuring the percentage of beaded axons as a function of time when cells are exposed to such drugs (to be published). When conducted on a sufficiently large sample size, this quantification can report on the neurotoxicity of therapeutic drugs. However, manual analysis of large datasets is not only laborious but also highly subjective, making comparisons across laboratories difficult.

There are many techniques and algorithms for biomedical image analysis and segmentation. They can be coarsely divided into classical techniques and AI-based techniques. While AI-based techniques are not far from infancy, classical methods are well established. Often, the choice depends on the requirements and trade-off between ease of usage and the effort needed to train AI. In analysing axonal morphology using neuronal culture, classical methods provide versatility and capability beyond mere segmentation. We often require precise measurements of various axonal metrics over time as the axon deteriorates. These include the average diameter of the axon, the evolution of axon diameter at the bead’s terminations, changes in the bead’s diameter and prominence, and the movement and merging of beads along the axon. As the study progresses, additional details will likely be required beyond those just mentioned here. This could involve unanticipated metrics or aspects of the axonal degradation process.

To address these challenges, we developed a software tool that tracks an axon through a time series of images. This Python-based tool identifies the onset of beading, provided a user selects the axon(s) in the initial frame using two or more mouse clicks. The code then measures the diameters of the selected axon at all pixels along its medial axis across all images in the series. This detailed frame-by-frame information provides crucial insights into the morphological evolution of beads, their movement along the axon, bead merging, and the spatial dependencies along the axon. These are important factors that can be used to compare normal axons with diseased axons or axons affected by chemical agents that are thought to cause neuronal damage. Hence, quantification of axonal shape changes using image processing algorithm is a powerful tool in investigating neurodegeneration. In the next few sections we discuss pre-processing of the images and the algorithm we have developed to obtain quantitative data on the time evolution of axonal beading and axonal diameter. Then we briefly discuss example experiments performed to induce axonal beading, and analysis of the results using the algorithm. Finally, we compare the efficacy and accuracy of the semi-automated analysis with human blind analysis. The entire code with comments, along with necessary details of parameters used for analysis, is available at https://github.com/vcpKumar/Axonal-Beading-Tracker. The repository also includes the necessary configuration files to adapt the algorithm to specific applications.

## 2 Computational pre-processing tools

### 2.1 Gaussian smoothing

Gaussian filters are often the preferred choice as low-pass filters. Filters like mean filters and median filters often have a preferred direction perpendicular to the edges [13]. Also filtering of the frequencies is not optimal, as will be evident from the Fourier transform of a box function, which is a *sinc* function with secondary peaks at higher frequencies introducing artifacts. Note that the mean filter is applied by convolving the image with a kernel of entries with an equal weight with the image to be smoothed.

The equation for a Gaussian filter can be written as 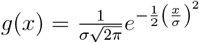. Smoothing is obtained by convolving the Gaussian with the image. In the frequency domain, Gaussian remains the same and convolution is replaced by multiplication. This gives a smooth cut-off for higher frequencies, unlike mean and mode filters. The Fourier transform of a Gaussian is given as 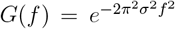. So frequencies that are filtered out are determined by the standard deviation *σ* of the Gaussian. To get a rough idea about the cut-off frequency, setting the exponent to 2 gives an attenuation of 1*/e*^2^, which is approximately 13.5%, giving a frequency value as *f* = 1*/πσ* cycles per pixel. In practice, when preparing an image for edge detection, *σ* is chosen such that the features of interest do not get washed out while random noise or speckles are smoothened out.

### 2.2 Hessian Analysis

Hessian analysis in image processing extracts the values of the principal curvatures at each pixel position. This is mainly used for feature extraction, particularly for ridge and valley detection. This technique is based on the Hessian matrix, which is a square matrix of the second-order partial derivatives of an image.

For an image *I*(*x, y*), the Hessian matrix *H* at a point (*x, y*) is given by:

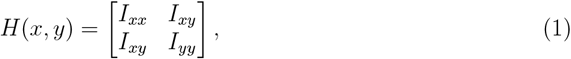

where *I*_*xx*_, *I*_*yy*_, and *I*_*xy*_ are the second-order partial derivatives of *I* with respect to *x* or *y* as indicated by the subscripts.

For an image, the Hessian matrix is a 2*×*2 matrix defined at each pixel position. So, at each pixel position, there will be two eigenvalues and eigenvectors, assuming that the matrices are non-singular. The highest eigenvalue gives the maximum value of the second derivative at a point, and the direction in which this occurs will be given by the corresponding eigenvector. Similarly, the other eigenvalue will represent the lowest value. At a minimum, the second derivatives are positive, while at a maximum, they are negative. In an image with variations in intensity (ups and downs), the ups or downs can be separated by choosing the right eigenvalues of the Hessian matrix [14, 15].

#### 2.2.1 Ridge / valley detection

Historically, the definition of ridges and valleys has been associated with landscape geometry. The same approach can be extended to the intensity landscape of images and it can be extended to multidimensional spaces also[16]. In biomedical images, for example, blood vessel and neurons appear as ridges. A simple thresholding can be a good option to segment such structures. However, in low contrast images, accurate detection of edges can be quite challenging. Several techniques have been described in the literature to enhance and segment vessel-like or neurite-like structures using Hessian analysis[17, 15, 18]. In our work we use the Meijering Filter to segment the axon of interest. The Meijering Filter uses a modified form of Hessian matrix given as [17, 19]

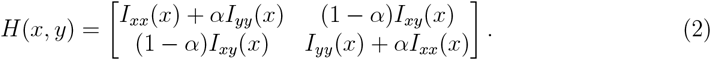

The authors have shown that choosing the value *α* = 1*/*3 gives optimum response for the filter along the length of the axon. To reduce noise, Gaussian smoothing is applied, prior to the Hessian analysis, facilitating a multiscale analysis [20, 21].

### 2.3 Wavelet based de-noising

While Gaussian smoothing does an excellent job in de-noising, it does not perform well in preserving edges and fine shape details well. The “washing out” of details causes random debris sticking to the axon or sharp protrusions on the axon to be confused with axonal beading when smoothed with a Gaussian kernel. So while Gaussian filtering is useful and efficient in identifying the broad outline of an axon, we do not rely on this method to identify beaded regions. Instead, we employ wavelet-based denoising, which effectively removes noise while preserving geometric features [22].

Wavelet analysis can be considered as a more generalized version of Fourier analysis. In Fourier analysis the resolution in conjugate domain would always be zero while the other domain will have a very high resolution which is set by the sampling rate. This comes with a disadvantage of not knowing where in space/time a particular frequency has occurred. Suppressing a frequency that has appeared only in a tiny window of the data will not be optimal in such cases. An immediate solution to this is to use a windowed Fourier Transform, where the sinusoidal waves are truncated by, say, a Gaussian window (Gabor Transform) or a rectangular window (spectrogram analysis) [23, 24]. A more robust strategy is the multi-resolution analysis using wavelets [23].

## 3 Description of the algorithm

### 3.1 Definition of key variables and libraries used

- `mask` -a binary mask which marks the likely contour of the axon. In an iteration, this will be based on the detected contour of the axon in the previous frame.
- `axonBin` -a binarized version of the axon under consideration.
- `spine` -the medial axis of the long tubular structure represented by an axon.
- `edges`-the two-pixel-thick boundaries of the tubular structure on either side.
- `maskFine` -a binary mask surrounding the exact location of the axon in a frame.
- `slps` -a vector containing the values of the local slopes of the axon.
- `highSlps` -a list containing the instances of slope values above a certain threshold.
- `beadCrop` -a window cropped around a possible bead after a preliminary analysis
- `beadBin` -a binarized image of `beadCrop`
- `beadMask` -a corresponding cropped image from `maskFine`
- `width` -width of a peak (FWHM) in the diameter plot
- `widthMax` -width of a peak at the base

The major libraries used in this work are NumPy [25], scikit-image [26], Insight Toolkit (ITK) [27], OpenCV [28], SciPy [29], pandas [30], PyWavelets [31], Numba [32], and Bokeh [33].

### 3.2 A walk through the algorithm

Figure 1 shows a flow-chart description of the algorithm. The program gets initialized by the user selecting an axon by clicking at its end points if it is straight, or at multiple points if the axon is curved. Figure 2a shows a curved axon selected by multiple mouse clicks. In some cases, the axonal position shifts as the axon evolves under the action of induced pharmacological perturbations. As the position does not change drastically across consecutive frames, we assume that the contour of the axon of interest will be enclosed within a region obtained by dilating its position in the previous frame, say, by 20 pixels or roughly 5× the typical axon diameter. As end points may also shift, the endpoints are identified in each frame using template matching. For this, windows of a fixed size (e.g., 40×40 pixels) around the axon’s endpoints in the previous frame are used as templates. Matching is performed over a larger window (e.g., 80×80 pixels) cropped from the current frame using the information from the previous frame. Correlation analysis is then performed, allowing for rotations and translations of the endpoint templates. In the code, this is executed by the function `templMatchMultiAng`

**Figure 1:**
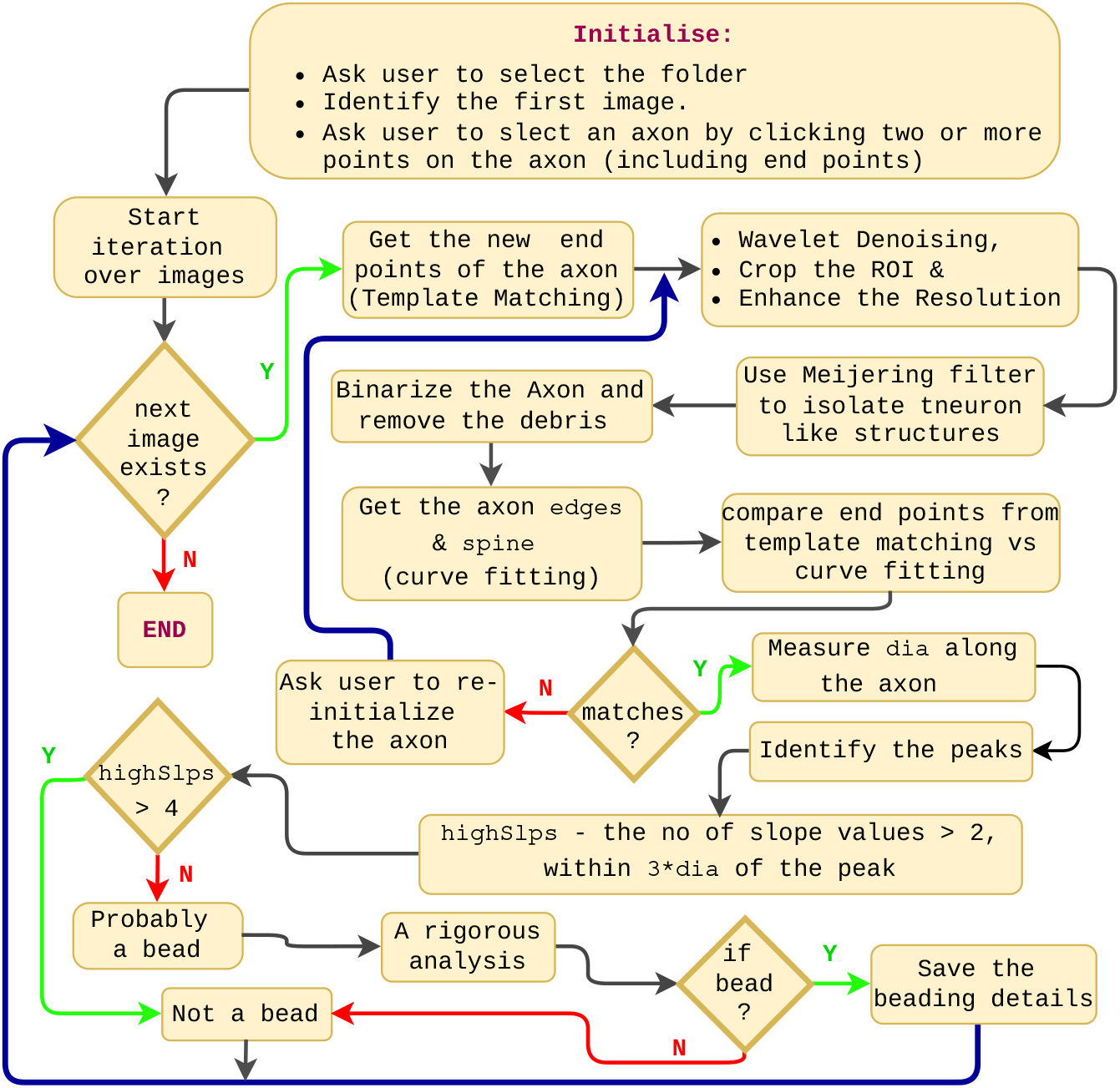
An overall description of the software pipeline.

**Figure 2:**
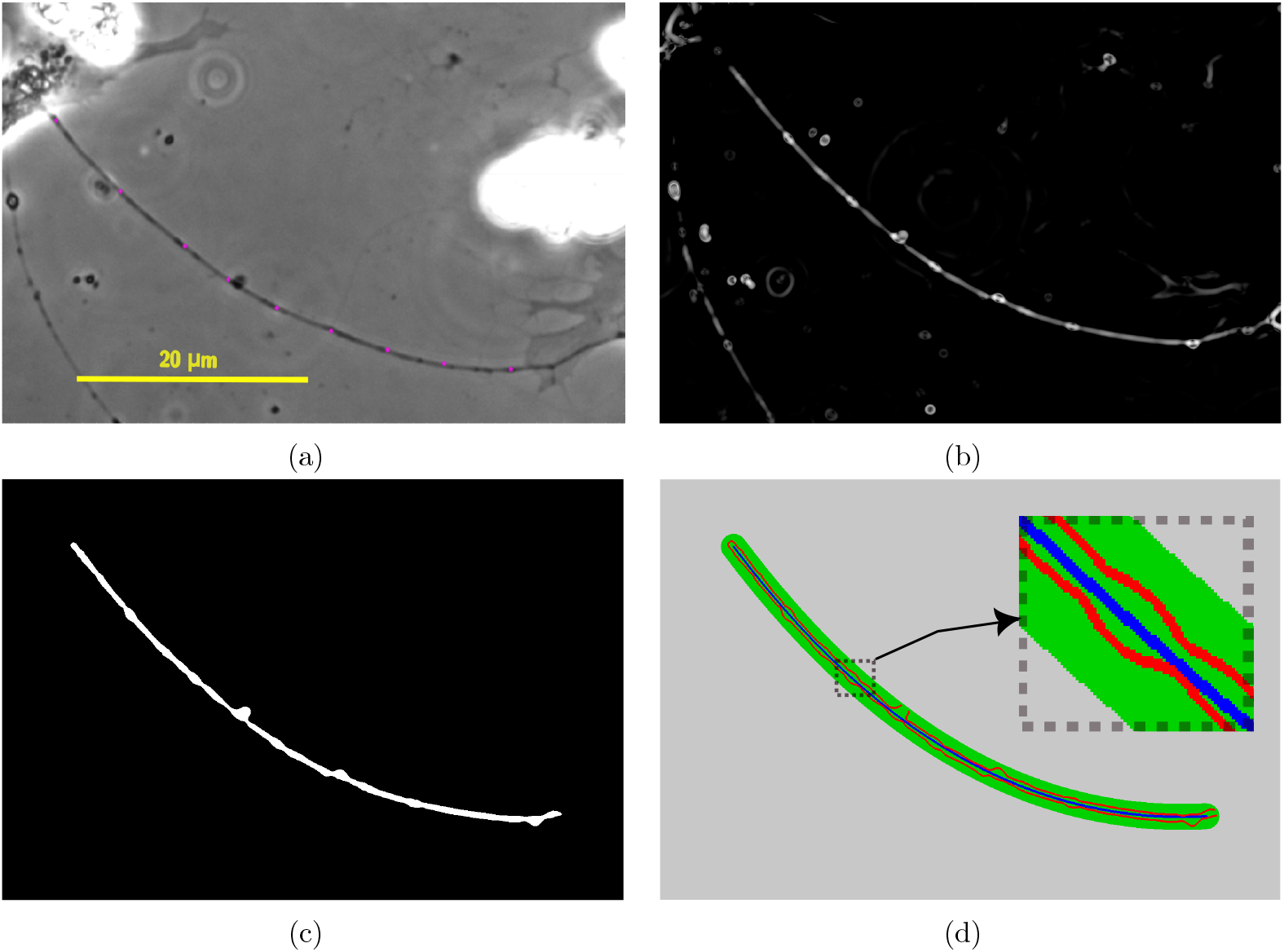
(a) Phase contrast image of an axon with points selected by the user via mouse-clicks shown as pink dots. (b) An image of the same axon taken after diameter modulations have set in (25^th^ frame), subjected to contrast enhancement using Meijering filter. (c) The binarized image of the same axon. (d) Image showing the detected edges and spine of the axon. The green background is the region selected by `maskFine`.

Wavelet denoising is performed using the function `denoise wavelet` from the python library `scikit-image`. A portion of the image containing the axon is cropped from the original image to reduce computational load. As the images were captured with a magnification of 20× on a camera with 1388×1040 pixels of size 6.45 *μ*m, the axon’s width, which is *∼* 0.5 *μ*m, would often be less than 5 pixels. So we enhance the resolution by a factor of three using bicubic interpolation [34] using the `OpenCV` function `cv.INTER_CUBIC`. Hessian analysis is employed to enhance the contrast of the axon. As discussed in section 2.2.1, Meijering filter is a robust implementation of the same, where tubular structures are given an enhancement based on the eigenvalues and eigenvectors of the Hessian matrix. With the Meijering filter, we look for axons at two different scales, *σ* = 3 and 6. Figure 2b shows an example of an axon after wavelet filtering and contrast enhancement for tubular structures. Note that this image corresponds to the 25^th^ frame of the axon shown in fig. 2a, after it has undergone beading (formation of swellings) along the axon, in response to treatment using the chemotherapeutic drug Vincristine, which causes neurodegeneration as a side effect.

Now, with the enhanced image at hand, it can now be binarized using a suitable threshold. The fact that the choice of the threshold can affect the width of the axon is not a matter of concern here, as this preliminary analysis will be followed by a more rigorous analysis. Now, continuing with the current discussion, there will be other axons and cellular structures in addition to the one we selected. To remove this the `mask` of the current frame, (`maskFine` from the previous image) is employed. This gives a more refined binary image of the axon. With the aid of morphological filtering, we can further refine it and the result obtained for the axon under consideration is shown in fig. 2c. In the code, this is executed by the function `morphFilt`. The exact sequence of operation can be found in the code. Skeletonizing `axonBin` and fitting a third-order polynomial to it yields the `spine` (midline) of the axon. This will be used later in finding the diameter of the axon. Two-pixel-thick `edges`, for diameter measurement, can be obtained by subtracting `axonBin` from its two-pixel-dilated version. This is shown in fig 2d, where the red lines shows the edges detected, the blue line the `spine`, and the green region shows the `maskFine` of the current frame. Note that this will be used as the `mask` for the subsequent frame.

The next step is to determine the diameter of the axon at each pixel position along the `spine`. The algorithm for the same is described in section 3.3. With the diameter at each pixel position determined, the peaks in the diameter values are indicative of beading. To further refine the analysis, suspected beaded regions of the axon are cropped out and a rigorous analysis is performed to confirm the presence of beading. The algorithm for the same is described in section 3.4

Upon completion, the analysis generates a text file containing beading data for each frame.We have set the condition for the onset of axonal beading as the presence of three beads in four continuous frames. Here, the time corresponding to the first of the four frames will be noted as the onset of beading.

### 3.3 Measurement of diameter and detection of beads

The algorithm for measuring diameter is illustrated in fig. 3. In the code, this is executed by the function `calcAxonDia`. Inputs to the algorithm are `slopes`, `edges`, and `spine`. The latter two are already available, and the equation for the slopes is derived from the analytical differentiation of the equation representing the `spine` (polynomial fit). By substituting the pixel coordinates into this derived equation, the slope values at each point along the `spine` are calculated. Maintaining `edges` at two-pixel thickness addresses the issue of 8-connected pixels when identifying overlapping pixels between two lines.

**Figure 3:**
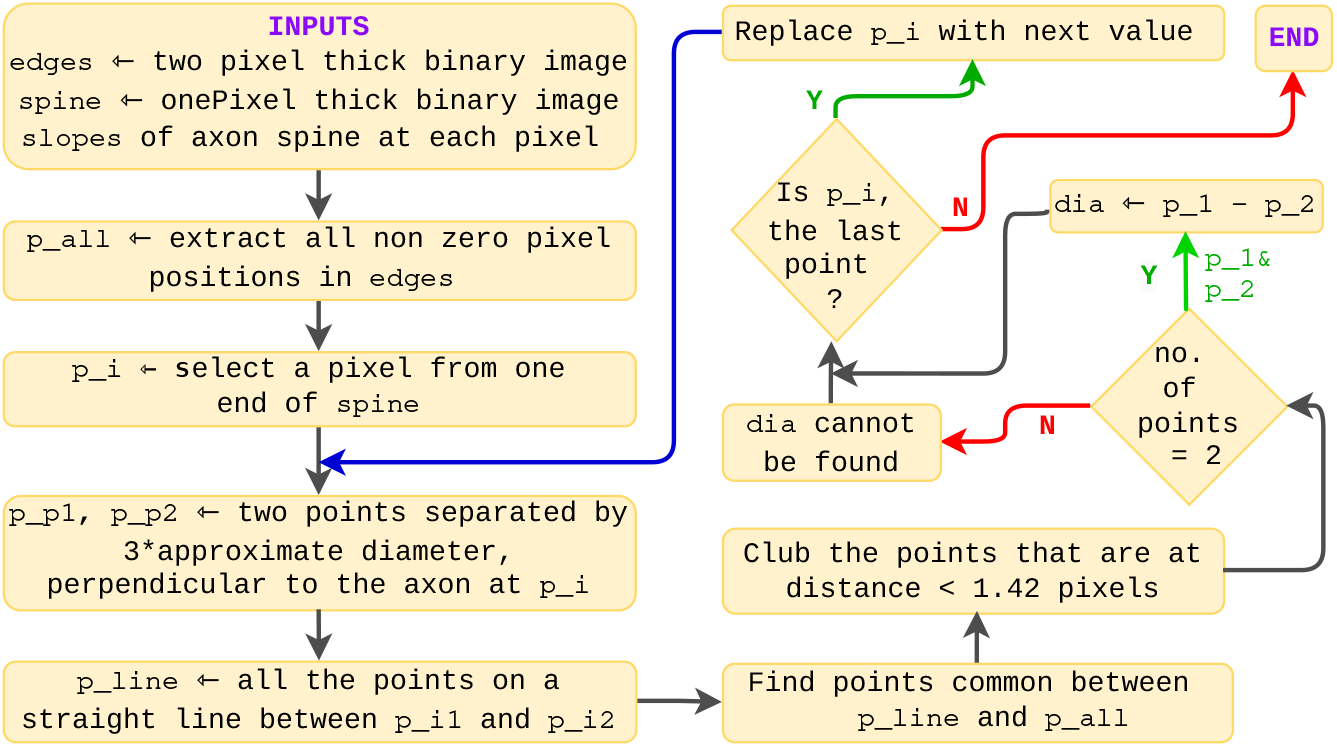
Algorithm for diameter measurement

Now, if the slope the axon at some pixel location *i* is *s*_*i*_, then the slope of the perpendicular to the axon at that point is *s*_*i⊥*_ = *−* 1*/s*_*i*_. Now a unit vector in the perpendicular direction can be written as

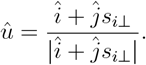

Knowing the unit vector, two points perpendicular to the axon, on either side, can be found. All the pixel coordinates connecting these two points are calculated using the Bresenham’s line algorithm [35].

The diameter of the axon is calculated as the distance between the intersecting points of the perpendicular with the `edges`. This can be found by extracting the coordinates of non-zero pixels from edges and comparing them with the pixel coordinates of the perpendicular line. Due to pixelation, there will be more than two points. This can be reduced to two points by clubbing points that are separated by a distance of less than 1.42 pixel. The algorithm for clubbing is shown in the fig. 1 of the supplementary data Note that the standard deviation of the Gaussian filter and value chosen for thresholding can have an effect on the detected diameter. Figure 4a shows a typical measurement procedure. Also, this is better demonstrated in the video 1 of the supplementary material. This is obtained using the 25^th^ frame for the same axon shown in fig. 2, where beading has occurred. Detected perpendicular lines are shown at 30-pixel intervals along the spine in fig. 4a for clarity.

**Figure 4:**
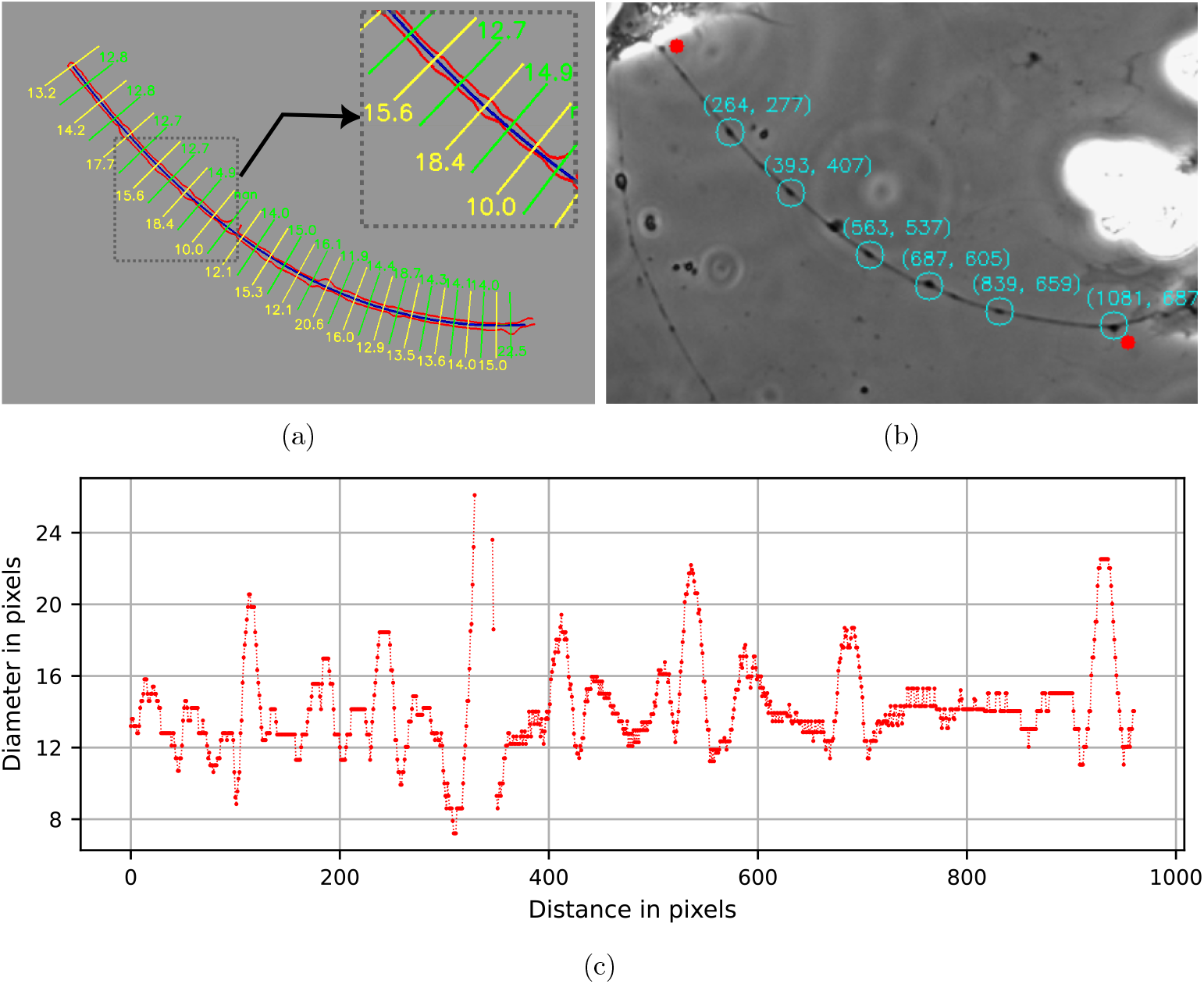
(a) Image showing the diameter detection procedure. The perpendiculars are shown only for every 30 pixels for clarity. (b) Image showing the beads that were identified using the peaks in the diameter plots. The red dots are the end points identified by template matching. (c) Plot of the diameter values along the axon length

Once the diameter along the axon is obtained, the peaks in the diameter are indicative of beading. In order to avoid cellular debris that may attach to the axons and be confused with beading, we apply an additional criterion. As beading is characterized by a smooth variation of slope along the axon, any shape with absolute values of slope higher than a threshold is considered as a debris. Figure 4c shows the diameter values plotted along the axon spine. The peaks are identified using the command `find_peaks` from the python library `scipy.signal`. We used prominence as a criterion to identify a peak. Prominence specifies how well a peak is distinguished from its surroundings [36]. We find that a good criterion for a peak to be a potential bead is that *prominence* = (*d/*4) where *d*, is the “representative diameter” of the axon, which is determined based on the mode of diameter values, after rounding them to whole numbers. Once the mode is identified, t he a verage d iameter is calculated by considering only those values that fall within the range of *mode* ± 2. To be considered as a possible bead, it is also required that the `highSlps` *<* 4, where the threshold for `highSlps` is set at 4. As, this is only a preliminary screening, the constraints are rather relaxed here. Figure 4b shows the location of the possible beads identified by t he c ode at this stage.

Note that the diameter measurement is a computationally intensive task. For an axon 1000 pixel long (after pixel extrapolation), perpendiculars has to be drawn at every point and this has to be repeated over the entire series of images. So the code for diameter measurement is written in Numba a python library that can render speeds comparable to *C* or FORTRAN using the LLVM compiler [32]

The peaks identified, at this stage, are on the reconstructed image given by the Meijering filter, which enhances tubular structures. And this, along with the inherent Gaussian smoothing involved, can cause small debris and protrusion on the axon to look like a bead, *ie*, the slope around them can become much less. To overcome this, knowing the position of a possible beads, a rigorous analysis is performed as discussed below.

### 3.4 Rigorous analysis to confirm beading

The detailed algorithm for rigorous analysis is described in fig.5. The function `rigBeadAnal` executes this in the code. For this, a 90×90 pixel window, `beadCrop`, centered at the suspected beading location, is cropped from the wavelet-filtered image. Figure 6a shows an example of the same. The other inputs include`beadBin` (corresponding cropped window from `axonBin`), `beadMask` (corresponding cropped window from `maskFine`), and the average diameter of the axon.

**Figure 5:**
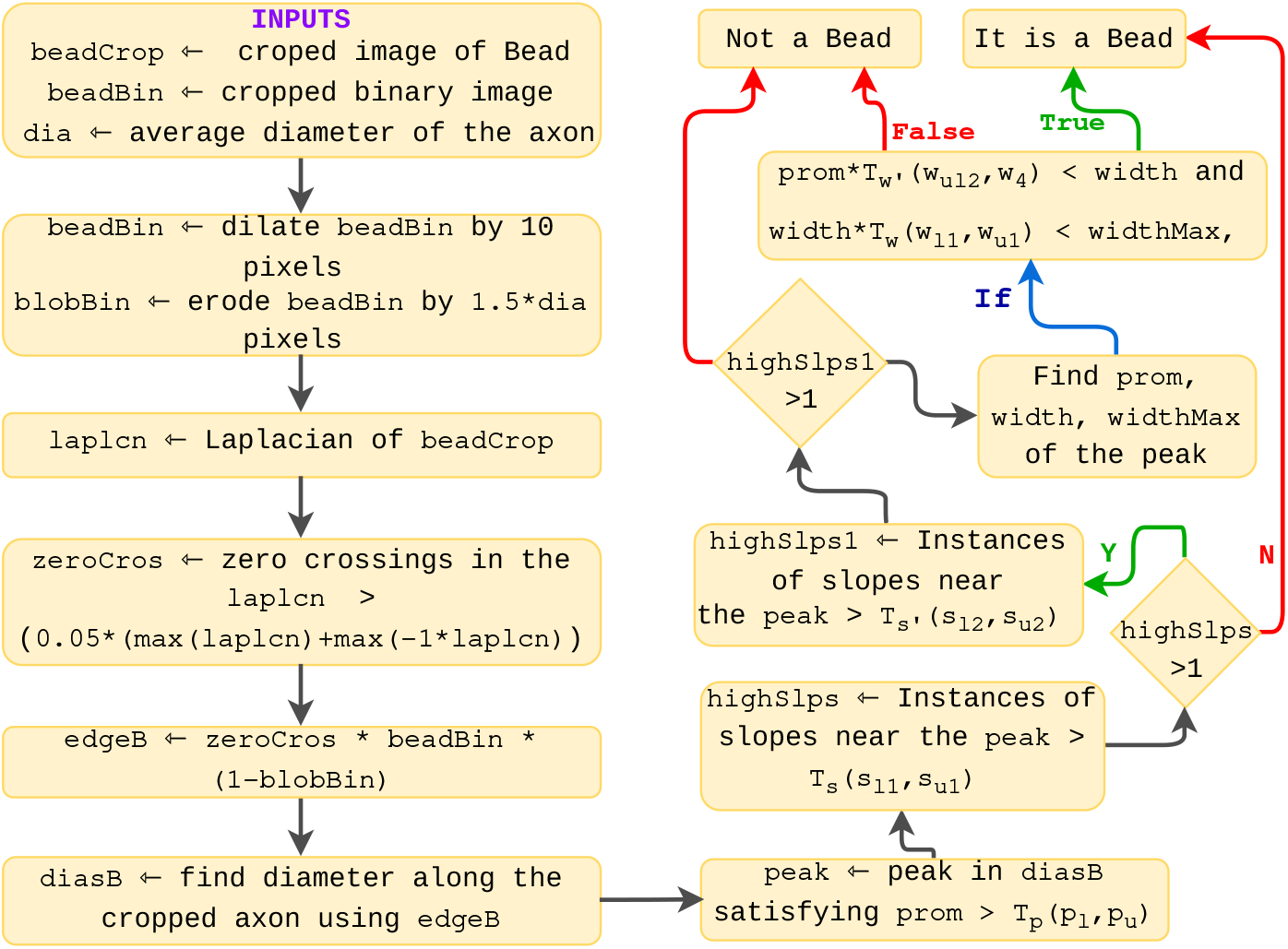
Algorithm for rigorous bead analysis

**Figure 6:**
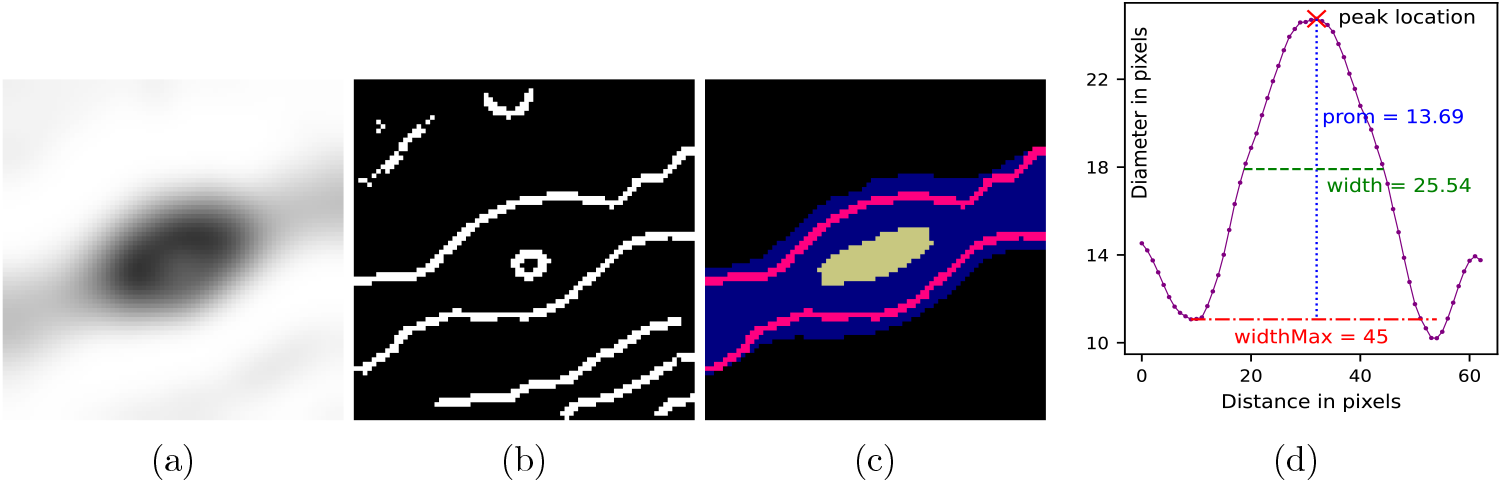
(a) Cropped image containing a suspected bead. (b) Edges traced from the Laplacian using zero-crossing. (c) Pink line shows the identified edges of the beaded region, obtained using the masks shown in blue and yellow. (d) Plot of the axonal diameter showing and the properties of the peak used for rigorous analysis.

In phase-contrast images, beading appears as a dark region surrounded by a white halo. At points of inflection, where there is a transition from dark to bright, the Laplacian of the image will give zeros or close to zero, with a sharp transition across zero. We consider these points of inflection as the axonal edges, which are identified by detecting zero-crossings in the Laplacian of the image. Two thresholds, *t*_1_ and *t*_2_ are used to determine the acceptable zero crossings.

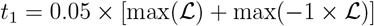

and

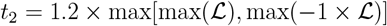

where ℒ is the Laplacian. The first condition defines a threshold for an acceptable zero-crossing in the Laplacian, while the second ensures a minimum contribution toward the threshold from both sides of the zero-crossing.

This method makes the edge detection immune to intensity fluctuations, as the calculation depends upon second derivative of intensity variation. Conventional edges detectors like Canny will detect multiple edges as there are large gradients occurring in different places and directions when beading occurs. For example, there will be a large gradient when you move from the centre of the bead to any direction. This often results in Canny generating an elliptical edge around the bead.

Figure 6a shows a cropped region of an axon containing a bead. It is quite obvious that there are multiple zero crossings in the image. This can be clearly seen in fig. 6b, where zero-crossings in the Laplacian are detected. A central circle can also be observed inside the bead when it becomes thicker. This occurs because phase contrast images are inherently an interferogram, where the intensity is a function of modulo 2*π*.

To remove irrelevant edges, the code generates two masks: the first by dilating `beadBin` by 5 pixels, and the second by isolating the very thick beaded regions with a “white top-hat” operation [37] and then by eroding it by seven pixels. The zero crossings are multiplied by these two masks to give the edges of the bead. This can be seen in fig. 6c, where the blue and yellow regions corresponds to the two masks respectively. With this a precise outline of the bead is obtained as shown in red. Now the diameter can be measured as described in section 3.3 and peaks in the diameter can be analyzed and confirmed to be a bead or not.

Figure 6d shows the plot of the measured diameters along the length of the axon. For the largest peak in the plot, the following parameters are computed: `prom` — the prominence, `width` — the full width at half maximum (FWHM), `widthMax` — the width at the base of the peak, and the gradient of diameter values within `widthMax` of the peak. A peak is classified as originating from a bead by thresholding these parameters. However, there are no single values for any of these parameters that suit all axons with varying diameters. To address this, these thresholds are set using a modified tanh function, as given below:

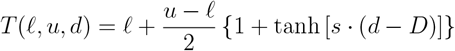

where,

‘*T* (*ℓ, u, d*)’ is the adaptive threshold to be found for the average diameter ‘*d*’, ‘*ℓ*’ is the lower bound of the threshold,

‘*u*’ is the upper bound of the threshold,

‘*s*’ is a scaling factor that controls the steepness of the transition from ‘*ℓ*’ to ‘*u*’, and

‘*D*’ is the mean value of the diameter for a population of unperturbed axons.

To further explain, consider the case of setting a threshold for prominence for an axon of diameter *d*. For this, we calculate *T*_*p*_(*p*_*ℓ*_, *p*_*u*_, *d*), which is the prominence threshold calculated as a function of diameter with lower and upper bounds set by *p*_*ℓ*_ and *p*_*u*_. Empirical observations suggest that the threshold should range from 4.5 to 6.5 pixels, corresponding to thin and thick axons, respectively. So we set *p*_*ℓ*_ = 4.5, *p*_*u*_ = 6.5, *s* = 1, *d*, and *D* = 15.5 (a central value for axon diameters, obtained empirically, as axon diameters typically vary between 13 and 18 pixels).

Next, the threshold for slope is set as *T*_*s*_(*s*_*ℓ*1_, *s*_*u*1_), with typical values of *s*_*l*1_ = 1.2 and *s*_*u*1_ = 0.9. If a peak satisfies *T*_*s*_, it is immediately classified as a bead. While large slope values *>* 1, are generally not indicative of beading, large slopes around 1.7 are not uncommon. To address this, an additional threshold *T*_*s*_*′* (*s*_*ℓ*2_, *s*_*u*2_) is introduced, with typical values *s*_*ℓ*2_ = 1.8 and *s*_*u*2_ = 1.3. If a peak satisfies only *T*_*s*_*′*, it must meet two additional criteria to be classified as a bead:

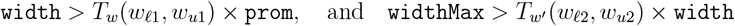

Typical values used for *w*_*ℓ*1_, *w*_*u*1_, *w*_*ℓ*2_, and *w*_*u*2_ are 1.3, 1.7, 1.5, and 1.9, respectively. This rigorous analysis requires reasonably good image quality. Otherwise, if the *T*_*s*_*′* criterion is satisfied, the peak is assigned to a bead without further checks.

Additionally, we analyze only one peak in the cropped regions. This is justified as we are primarily looking for the onset of beading, where it is highly unlikely for two beads to appear in the same cropped window. However, in heavily beaded axons, multiple beads in a single window are possible. If necessary, such cases can be handled by setting a height threshold for the peaks relative to the axon diameter.

## 4 Neuronal cell culture and induction of axonal beading

Neuronal cultures were obtained from 9 day old chick embryos as detailed in [7]. Sensory neurons from the Dorsal Root Ganglia were grown on plane glass coverslips and imaged using a Phase Contrast microscope (Zeiss Axio Observer Z1) using a 20× objective. Timelapse images were recorded with a depth of 12 bits using a CCD camera with 1388×1040 pixels, each of size 6.45 *μ*m (Zeiss Axiocam MRm). In order to induce a morphological transformation, cells grown for 48 hr in culture were treated with the chemotherapeutic drug Vincristine at different concentrations and then time lapse images were acquired at a frame rate of one frame per minute. Like several chemotherapeutic agents, this drug induces neuropathy in patients as a side effect to cancer treatment. In cell culture, the drug causes extensive beading–appearance of swellings along the normally cylindrical axons. Susceptibility of axons to undergo beading may be used as an assay to screen drugs for the potential to cause unwanted neuronal damage.

## 5 Comparison between semi-automated and manual analysis

Vincristine, a widely used chemotherapeutic drug, is known for its neurotoxic side effects [12]. To assess the code’s effectiveness in detecting beading, we analyzed axonal images treated with Vincristine at concentrations of 100, 500, and 3000 nM. Following the standard experimental protocol, images were captured every minute over a 90-minute period.

Figure 7 compares the analysis performed by a human with that of the software for different Vincristine concentrations. As can be seen, there is good agreement between both methods, although some variation is expected due to inherent human bias. Additionally, the software was able to analyze about 20% fewer axons due to issues such as axons moving significantly between frames or becoming out of focus as time progresses. This drawback can be easily overcome by feeding images from more trials to the code.

**Figure 7:**
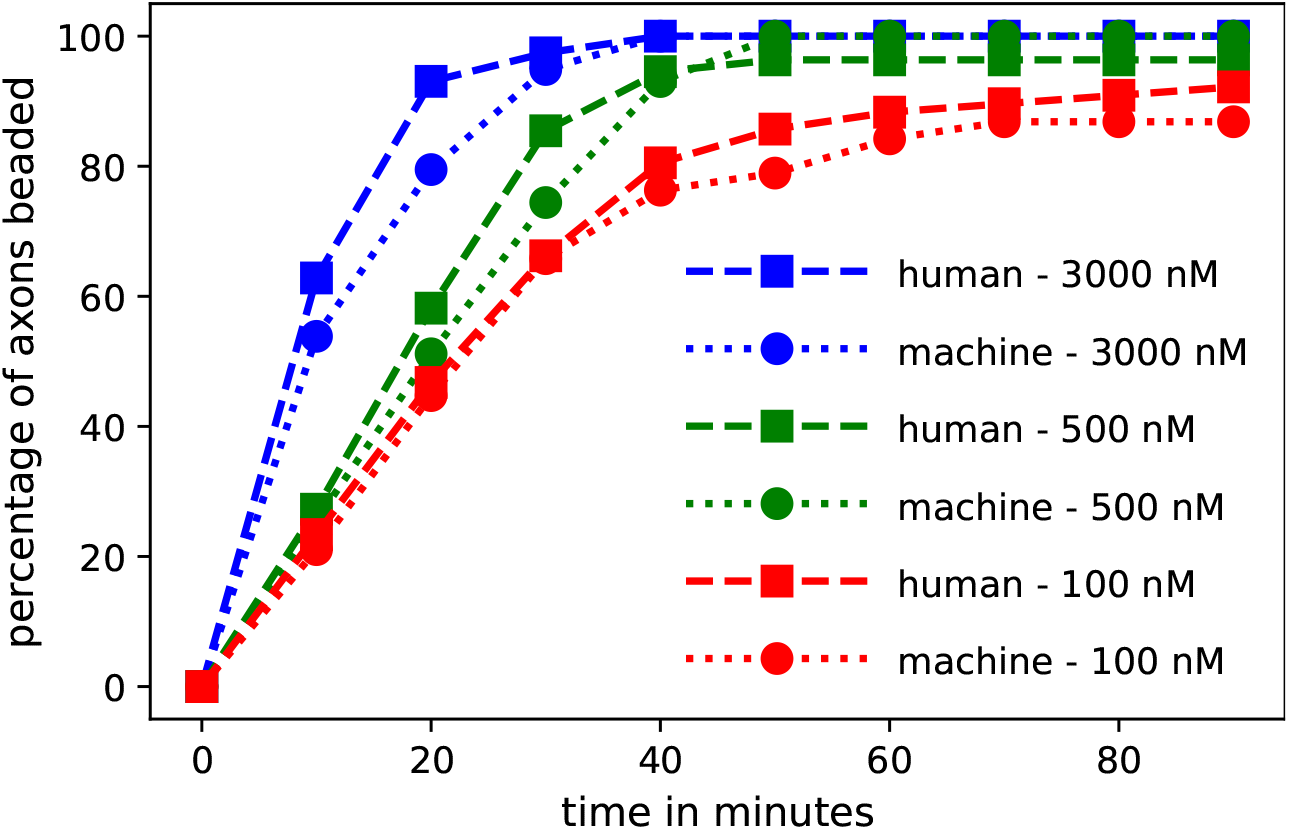
Comparison of beading analysis performed by human and software for various concentrations of the drug Vincristine

All analyzes were performed on a laptop with 8GB of RAM and an Intel^®^ Core™ i5-10210U processor, a quad-core CPU designed for light computational tasks. The processing time for analyzing an axon across 90 frames was approximately one minute. This performance could be significantly improved using a workstation, as matrix operations are CPU-intensive, and laptops often throttle performance due to thermal constraints. Additionally, capturing images at 40× magnification or camera with higher pixel density would reduce the need for up-scaling and simplify axon segmentation. High-contrast images like those that can be obtained using fluorescent dyes will also facilitate easier separation of the axon from the background. In conclusion, this semi-automated approach will greatly facilitate investigations on axon degeneration, especially that caused by pharmaceutical agents which causes axonal beading.

## 6 Discussion

The field of digital image processing encompasses a broad array of mathematical techniques and algorithms for various tasks such as segmentation and feature detection [38, 39, 40, 41], image registration [42, 43], 3D reconstruction [44], deconvolution [45, 46], and super-resolution techniques to overcome diffraction limits [47, 48]. With the advent of AI and deep learning, computational tools for addressing these aspects have significantly advanced, providing powerful new methods for tackling these challenges [49, 50].

The study of axonal morphology in a quantitative manner as a way to assess neurodegeneration under induced conditions is still in its initial stages and there are not many codes available for automated analyses. One of the few works we could find that has some similarity to our work is by Kilinc *et al*. [51]. One of the main tasks of their work was to quantify the extent of beading, for which they introduced a “beading score”. This score is calculated as directly proportional to the cube of the bead radius times the number of beads, and inversely proportional to the length of the axon. Their image analysis method involves many interactive subroutines for cleaning up, filling gaps and analyzing a single image, where users must manually remove, add, or connect pixels. Hence, it is not practical to analyze the temporal evolution of axonal beading for a large number of axons.

Furthermore, their radius measurement algorithm, which places disks of various diameters on the binarized axon and assigns the radius of the disk that overlaps by at least 80% as the axon’s radius at that position, is computationally efficient but may not be the best choice when high accuracy is prioritized. In contrast, our algorithm involves automated detection of perpendiculars at each point along the detected axon spine and searching for intersections on the axon skeleton, which can provide sub-pixel accuracies. Additionally, our use of Python, as opposed to MATLAB, should make our method more accessible to the wider research and development community.

Another interesting work is the automated detection of axonal synapses or boutons in 3-D two-photon images, without tracking the axons [52]. The researchers utilized the Laplacian of Gaussian (LoG) filter to highlight bouton-like structures with a suitable value of the standard deviation of the Gaussian. To distinguish true boutons from false positives, SVM (Support Vector Machine) was employed. The paper reports that after training SVM with a good set of extracted features, the algorithm achieved a recall rate of 95%. We do not resort to such methods as beading should be looked upon a a morphological change more than an intensity change. We are interested in detecting beads per axon and this requires automated tracing of axons as discussed in earlier sections. Furthermore, our approach allows for tracking of beads over time which is important when investigating the dynamics of beads during axonal atrophy induced by disease or drug treatment.

## 7 Conclusions

In this study, we developed a semi-automated software tool to efficiently analyze axonal beading, a morphological change indicative of axonal degeneration. The software automates axon tracking and beading detection using time series images.

This algorithm overcomes many of the limitations of previously reported ones. In particular, it enables efficient and accurate analysis of temporal evolution of beading in large datasets. The latter advantage is significant, as the axonal morphological changes caused by drugs are indicative of their potential to cause neuronal damage as a side effect, and hence beading assays can be used to investigate their neuropathy potential. Our results show strong agreement between software-based and manual analysis for various concentrations of the neuropathy-inducing chemotherapy drug Vincristine.

The tool performed well on a standard laptop, missing only 10% to 20% of axons compared to expert analysis.. Despite this minor limitation, the code offers an efficient and reproducible solution for large scale axonal beading studies. Further work could extend the code’s capabilities to address additional metrics relevant to axonal degeneration and apply it to different experimental conditions.

## Acknowledgments

We thank Serene Rose David and Neha Mohamad for the experiments and data. The authors acknowledge support from *The Wellcome Trust-DBT India Alliance* (Grant IA/TSG/20/1/ 600137).

