## Supplementary figures and images for "Semi-automated analysis of beading in degenerating axons"

### fig. 1

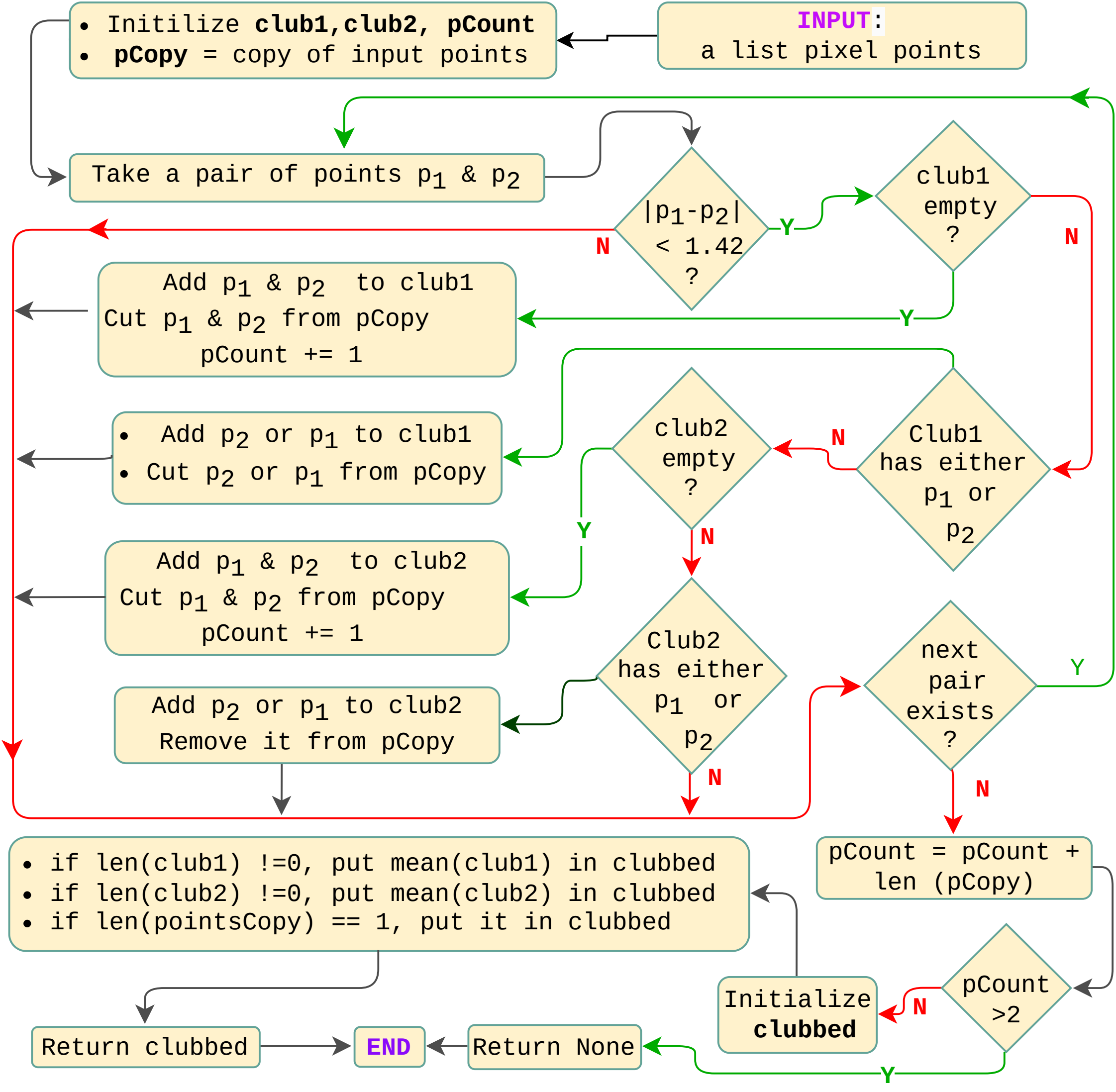
